## Supplementary Information for "Tunable genetic devices through simultaneous control of transcription and translation"

| **Supplementary Notes** | **Page** |
| --- | --- |
| Supplementary Note 1: Modelling the tunable expression system | 2 |
| Supplementary Note 2: Modelling possible retroactivity effects | 4 |
| **Supplementary Figures** | |
| Supplementary Figure 1: Characterization of sensor systems | 6 |
| Supplementary Figure 2: Simulated response functions for models | 7 |
| Supplementary Figure 3: Plasmid maps | 8 |
| Supplementary Figure 4: Flow cytometry automated gating approach | 9 |
| **Supplementary Tables** | |
| Supplementary Table 1: Modelling parameters | 10 |
| Supplementary Table 2: Sequences used for thermodynamic modeling | 11 |
| Supplementary Table 3: Performance summary of tunable NOR gate | 12 |
| Supplementary Table 4: Synthesized DNA fragments used in this study | 13 |
| Supplementary Table 5: Primers used in this study | 14 |
| **Supplementary References** | 15 |

**Supplementary Note 1: Modelling the tunable expression system**

To better understand the behavior of the tunable expression system (TES), we derived the following set of pseudo-chemical reaction equations to capture changes in the concentrations of the four main species in our system: toehold switch transcript *S*, tuner sRNA *T*, switch-sRNA complex *C*, and output protein *P* (see **Figure 1a** for a schematic of the device)

$$\xrightarrow{\alpha_T+\alpha_T^0} T\ , \tag{1}$$

$$\xrightarrow{\alpha_S+\alpha_S^0} S\ , \tag{2}$$

$$T + S \xrightarrow{k_{C+}} C\ , \tag{3}$$

$$C \xrightarrow{k_{C-}} T + S\ , \tag{4}$$

$$C \xrightarrow{\alpha_P} C + P\ , \tag{5}$$

$$P \xrightarrow{\gamma_P} \emptyset\ , \tag{6}$$

$$T \xrightarrow{\gamma_T} \emptyset\ , \tag{7}$$

$$S \xrightarrow{\gamma_S} \emptyset\ , \tag{8}$$

$$C \xrightarrow{\gamma_C} \emptyset\ . \tag{9}$$

Here, $\alpha_S$ and $\alpha_T$ are the induced promoter activities (rate of transcript production) of the main and tuner inputs, respectively, and $\alpha_P$ is the production rate of the output protein. $\alpha_S^0$ and $\alpha_T^0$ are the basal rates of transcription of the main and the tuner input promoters, respectively. $k_{C+}$ and $k_{C-}$ are the binding and unbinding rates of the toehold switch transcript with tuner sRNA. $\gamma_S$, $\gamma_T$, $\gamma_C$, $\gamma_P$ are first-order degradation rates of the main toehold switch transcript, tuner sRNA, switch-sRNA complex, and output protein, respectively. Stochastic simulations of these chemical reaction equations were performed using COPASI (**Methods**) and a Systems Biology Markup Language (SBML) file capturing the model is available in **Supplementary Data 1**.

From these chemical reactions, we derived a set of ordinary differential equations (ODEs) to model the dynamics of the TES and assess how its function is affected by key design parameters. The model tracks changes in the concentrations of the main toehold switch transcript *S*, tuner sRNA *T*, switch-sRNA complex *C*, and output protein *P*. It consists of the following four coupled ODEs that capture production and loss of each species due to complex formation or degradation

$$\frac{dS}{dt} = \alpha_s^0 + \alpha_S - k_{C+}ST + k_{C-}C - \gamma_S S\ , \tag{10}$$

$$\frac{dT}{dt} = \alpha_T^0 + \alpha_T - k_{C+}ST + k_{C-}C - \gamma_T T\ , \tag{11}$$

$$\frac{dC}{dt} = k_{C+}ST - k_{C-}C - \gamma_C\, C, \tag{12}$$

$$\frac{dP}{dt} = \alpha_P C - \gamma_P P. \tag{13}$$

All species and rates match those in the chemical reactions captured by **Equations (1)–(9)**. To analyze the behavior of this system, numerical simulations of these ODEs were performed using Python (**Methods**) across a range of biologically realistic parameter values (**Figure 3**; **Supplementary Table 1**).

**Supplementary Note 2: Modelling possible retroactivity effects**

One way that retroactivity [1, 2] could arise in the TES is due to the demand that synthesis of an output protein (e.g. YFP) places on the shared ribosome pool of a cell [3]. If expression of the output protein is sufficiently burdensome, there would be significant drop in the free ribosomes available to synthesize core machinery such as RNA polymerase (RNAP) and thereby a reduction in the transcription rates of the TES. To better understand this effect, we modified our existing model (**Supplementary Note 1**) to incorporate a factor $r_e$ that captures the relative availability of host resources (i.e. RNAP)

$$\frac{dS}{dt} = (\alpha_S^0 + \alpha_S) r_e - k_{C+} ST + k_{C-} C - \gamma_S S \,, \quad (14)$$

$$\frac{dT}{dt} = (\alpha_T^0 + \alpha_T) r_e - k_{C+} ST + k_{C-} C - \gamma_T T \,, \quad (15)$$

$$\frac{dC}{dt} = k_{C+} ST - k_{C-} C - \gamma_C \, C. \quad (16)$$

To calculate $r_e$ we coupled these equations to a delay differential equation (DDE) model of ribosome allocation dynamics [3]. This model considers a fixed total pool of ribosomes $R_t$ and their allocation between endogenous transcripts $E$ (modelled as a single “average” type, see [3] for details), translationally active TES complexes $C$, and a free ribosome pool $R_f$. Conservation of the total number of ribosomes per cell is ensured by the additional algebraic constraint

$$R_f = R_t - [R_E + R_C(t)], \quad (17)$$

where $R_E$ and $R_C$ represent allocation of ribosomes to endogenous transcripts and active TES complexes, respectively, whose changes are captured by the following equations

$$\frac{dR_E}{dt} = \alpha_E R_f(t) E - \alpha_E R_f(t - \tau_E) E, \quad (18)$$

$$\frac{dR_C}{dt} = \alpha_C R_f(t) C(t) - \alpha_C R_f(t - \tau_C) C(t - \tau_C). \quad (19)$$

Here, $\alpha_E$ and $\alpha_C$ are the translation initiation rate of the endogenous transcripts and active TES complexes, respectively. Each equation captures the initiation of free ribosomes onto a respective transcript and the removal of ribosome from a transcript back into the free ribosome pool after the specific translation time of the transcript, either $\tau_E$ for endogenous transcripts or $\tau_C$ for active TES complexes. The relative availability of core cellular machinery like RNAP is then given by

$$r_e = \frac{R_E}{R_0}, \quad (20)$$

where $R_0$ is the steady state value of $R_E$ in the model when only endogenous genes are present (i.e. no TES complexes are considered, $C$ = 0).

Analysis of the model was performed by numerical simulation of the DDEs above using Julia (**Methods**) across a range of biologically realistic parameter values (**Supplementary Table 1**). We then considered two cases: 1. when TES transcript synthesis rate was coupled to ribosome availability (as above), and 2. when there was no coupling between the output protein production rate and the synthesis rate of the TES transcripts (i.e. $r_e$ = 1). The simulations showed that retroactivity (case 1) could impact the response function of the TES device, but only when the output caused a significant burden on the cell and only for the most highly expressed outputs, i.e., when both the input and tuner promoter activities were high (**Supplementary Figure 2**).

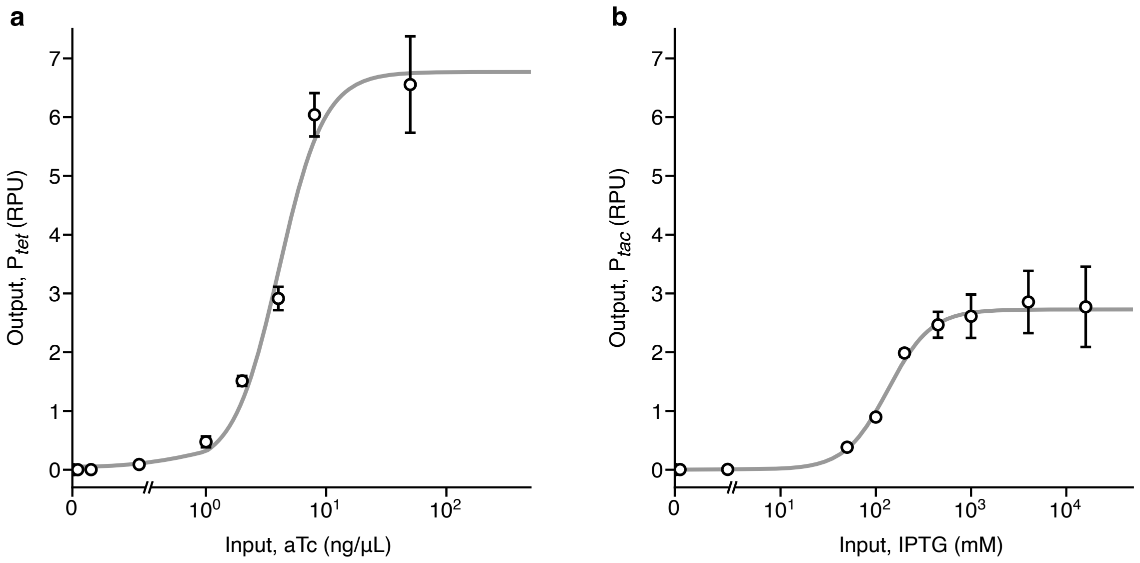


**Supplementary Figure 1: Characterization of sensor systems.** (**a**) Response function of the $P_{tet}$ promoter to varying aTc concentrations: 0, 0.01, 0.04, 0.14, 0.5, 1, 2, 4, 8, 50 ng/µL. (**b**) Response function of the $P_{tac}$ promoter to varying IPTG concentrations: 0, 0.5, 5, 50, 100, 200, 450, 1000, 4000, 16000 mM. Points denote average output in relative promoter units (RPU) (**Methods**) from three biological replicates with error bars showing ±1 standard deviation. Solid grey lines show fitted Hill functions ($P_{tet}$ fit parameters: $y_{\mathrm{min}}$ = 0.05 RPU, $y_{\mathrm{max}}$ = 6.77 RPU, $K$ = 4.0 ng/µL aTc, $n$ = 2.3; $P_{tac}$ fit parameters: $y_{\mathrm{min}}$ = 0.001 RPU, $y_{\mathrm{max}}$ = 2.73 RPU, $K$ = 134 mM IPTG, $n$ = 1.9).

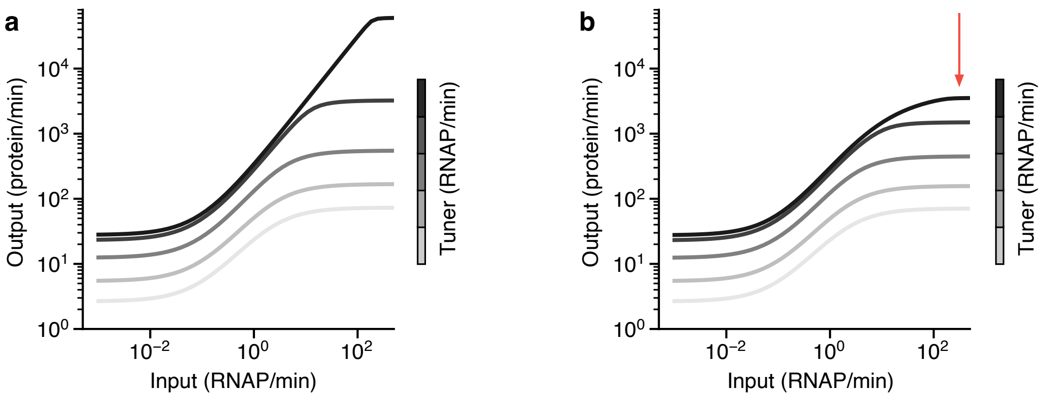


**Supplementary Figure 2: Simulated response functions for models.** (**a**) Response functions when TES transcription is not coupled to varying availability of core cellular resources ($r_e$ = 1). (**b**) Response functions when TES transcription is influenced by varying availability of core cellular resources due to synthesis of the output protein. Red arrow shows the decrease in the TES protein output due to retroactivity. All plots show individual curves for tuner promoter activities: 0.0001, 0.3, 1.5, 10, 190 transcripts/min (light grey to black). Model described in **Supplementary Note 2**.

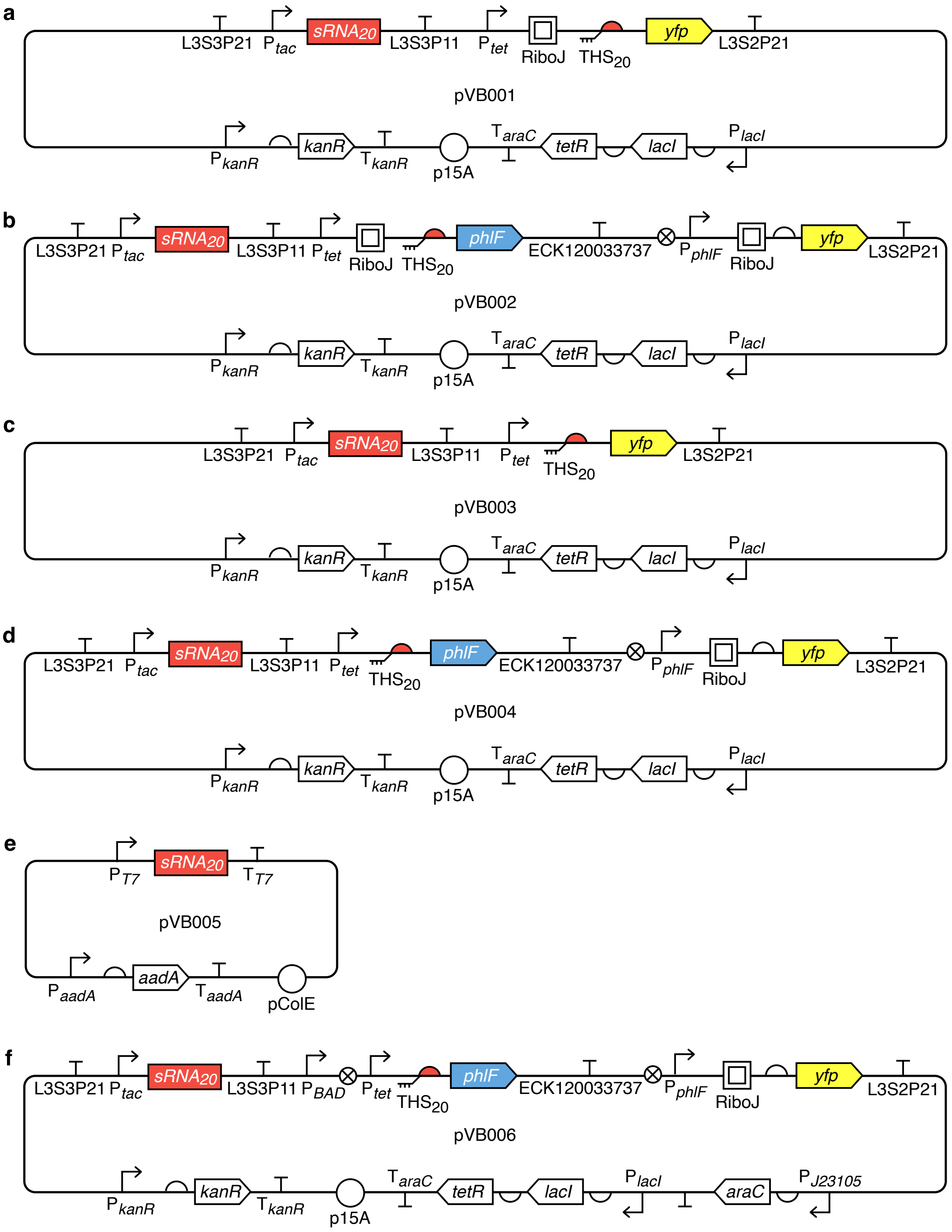


**Supplementary Figure 3: Plasmid maps.** (**a**) pVB001: Tunable expression system. (**b**) pVB002: Tunable genetic NOT gate. (**c**) pVB003: Tunable expression system without RiboJ insulating element. (**d**) pVB004: Tunable genetic NOT gate without RiboJ insulating element. (**e**) pVB005: Tuner sRNA booster. (**f**) pVB006: Tunable NOR gate.

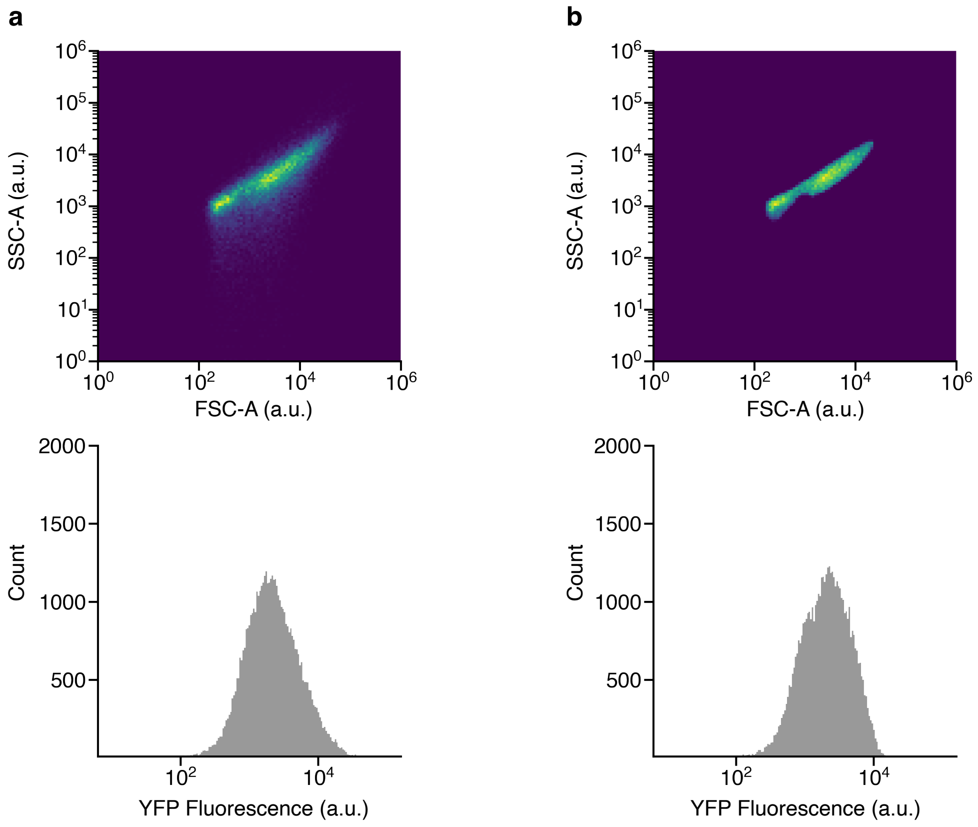


**Supplementary Figure 4: Flow cytometry automated gating approach.** (**a**) Raw event data from the flow cytometer. (**b**) Automatically gated event data with unwanted debris removed (**Methods**). Gating is performed using FlowCal which implements density-based filtering to discard events that deviate in their appearance from the main population. Top pseudo color density scatter plots show the forward and side scatter area (FSC-A and SSC-A) and bottom histogram shows the associated raw YFP fluorescence from these events. Data is collected for the initial TES construct with RiboJ present and an input of 6.6 RPU and tuner of 2.6 RPU.

**Supplementary Table 1: Modelling parameters.**

| Name | Description | Value(s) | Units | Refs. |
|---|---|---|---|---|
| $\alpha_S$ | Induced transcription rate of toehold switch promoter | 0–300[a] | transcripts min$^{-1}$ | [4,5,6] |
| $\alpha_T$ | Induced transcription rate of tuner sRNA promoter | 0–190[b] or 730[c] | transcripts min$^{-1}$ | [4,5,6] |
| $\alpha_S^0$ | Basal transcription rate of toehold switch promoter | 0.0886[d] | transcripts min$^{-1}$ | [4,5,6] |
| $\alpha_{T^0}$ | Basal transcription rate of tuner sRNA promoter | 0.2307 or 0.9228[d] | transcripts min$^{-1}$ | [4,5,6] |
| $\alpha_P$ | Protein synthesis rate from toehold switch | 5 | proteins complex$^{-1}$ min$^{-1}$ | [7] |
| $k_{C+}$ | Association rate of switch and trigger RNAs into complex | 0.0257[e] | complexes transcript$^{-1}$ min$^{-1}$ | [8] |
| $k_{C-}$ | Dissociation rate of switch and trigger RNAs from complex | 0.00672[e] | transcripts complex$^{-1}$ min$^{-1}$ | [8] |
| $\gamma_S$ | Degradation rate of toehold switch transcript | 0.231[f] | min$^{-1}$ | [9] |
| $\gamma_T$ | Degradation rate of trigger sRNA | 0.231[f] | min$^{-1}$ | [9] |
| $\gamma_C$ | Degradation rate of toehold switch and sRNA complex | 0.231[f] | min$^{-1}$ | [9] |
| $\gamma_P$ | Degradation rate of output protein | 0.035[g] | min$^{-1}$ | [10] |
| $R_t$ | Total number of ribosomes | 26300 | ribosomes | [3] |
| $E$ | Number of endogenous transcripts | 4140 | transcripts | [3] |
| $\alpha_E$ | Translation initiation rate of endogenous transcripts | 0.0002 | ribosome$^{-1}$min$^{-1}$ | [3] |
| $\alpha_C$ | Translation initiation rate of active TES complexes | 0.0034 | ribosome$^{-1}$min$^{-1}$ | [3] |
| $\tau_E$ | Translation time of average endogenous gene | 0.28 | min | [3] |
| $\tau_C$ | Translation time of TES output gene | 5.67 | min | [3] |

a. We assume that the induced translation initiation rate of $P_{tet}$ is similar to the transcription rate of an average constitutively expressed endogenous gene (20 transcripts min$^{-1}$) [2] and that a plasmid with a p15A origin of replication has 15 copies per cell [3].
b. The induced transcription initiation rate of $P_{tet}$ is 1.57 times higher than $P_{tac}$ [1].
c. We assume the pColE1 origin of replication leads to 50 copies per cell [7] and thus the transcription initiation rate from all $P_{tac}$ promoters is ~4 times higher when the sRNA booster is present.
d. Uninduced basal expression calculated based on fold-change activation measured in previous work [1] and the estimated induced transcription initiation rates ($\alpha_S$, $\alpha_T$).
e. We assume the rates of DNA hybridization are similar to RNA hybridization [5].
f. We assume the half-life of the switch transcript and tuner sRNA are similar to the average mRNA half-life in a cell, measured to be 3 min in exponentially growing cells [6].
g. Dilution by growth is the dominant mode of degradation. The doubling time of cells during these experiments was determined to be ~20 min.

**Supplementary Table 2: Sequences used for thermodynamic modeling.**

| Name | RNA sequence |
|---|---|
| Toehold Switch | GGGCGUUAAUCUCUGGCUUGCUUUAUGUCUGUAAACAGAGGAGAUACAGAAUGAAAGCAAGCAACCUGGCG GCAGCGCAAAAGAUGCGUAAA |
| Tuner sRNA | GGGUCAUGACUGGGACACGCCAGUCAUGAGAAUACAGACAUAAAGCAAGCCAGAGAUUAACGAAG |
| Cleaved Toehold Switch with RiboJ | UCACCGGAUGUGCUUUCCGGUCUGAUGAGUCCGUGAGGACGAAACAGCCUCUACAAAUAAUUUUGUUUAAG GGCGUUAAUCUCUGGCUUGCUUUAUGUCUGUAAACAGAGGAGAUACAGAAUGAAAGCAAGCAACCUGGCGG CAGCGCAAAAGAUGCGUAAA |
| Full Toehold Switch with RiboJ | AGCUGUCACCGGAUGUGCUUUCCGGUCUGAUGAGUCCGUGAGGACGAAACAGCCUCUACAAAUAAUUUUGU UUAAGGGCGUUAAUCUCUGGCUUGCUUUAUGUCUGUAAACAGAGGAGAUACAGAAUGAAAGCAAGCAACCU GGCGGCAGCGCAAAAGAUGCGUAAA |

**Supplementary Table 3: Performance summary of tunable NOR gate.**

| **Tuner** (RPU) | **Dynamic range**[a] (a.u.) Ara / aTc | | | **Fold-change**[a] Ara / aTc | | | **Intersection**[a] Ara / aTc | | |
|---|---|---|---|---|---|---|---|---|---|
| | + / – | – / + | + / + | + / – | – / + | + / + | + / – | – / + | + / + |
| 0.002 | 3210 ± 573 | 2623 ± 322 | 3671 ± 233 | 5.6 ± 3.9 | 4.3 ± 2.6 | 16.7 ± 3.3 | 0.21 ± 0.12 | 0.13 ± 0.07 | 0.04 ± 0.003 |
| 2.6 | 3808 ± 604 | 4013 ± 718 | 4012 ± 729 | 8.4 ± 0.1 | 15.3 ± 4.5 | 13.9 ± 4.5 | 0.07 ± 0.04 | 0.08 ± 0.02 | 0.05 ± 0.02 |

a. Measurements are a comparison to experiments when both input inducers (Ara and aTc) are absent. Average values are shown ± 1 standard deviation calculated from flow cytometry data for three biological replicates.

**Supplementary Table 4: Synthesized DNA fragments used in this study.**

| Name | Sequence |
|---|---|
| TES-P1 | CGCAAACCGCCTCTCCCCGCAACGATCGTTGGCTGTGTTGACAATTAATCATCGGCTCGTATAATGTG<br>TGGAATTGTGAGCGCTCACAATTGGGACCGTGGACCGCATGAGGTCCACGGTAAACATAACTATAACA<br>AGCCTACAATTCATTCAAACCCAATTATTGAACACCCTTCGGGGTGTTTTTTTGTTTCTGGTCTACCT<br>ACTCCACCGTTGGCTTTTTTCCCTATCAGTGATAGAGATTGACATCCCTATCAGTGATAGAGATAATG<br>AGCACAGCTGTCACCGGATGTGCTTTCCGGTCTGATGAGTCCGTGAGGACGAAACAGCCTCTACAAAT<br>AATTTTGTTTAAGGGCGTTAATCTCTGGCTTGCTTTATGTCTGTAAACAGAGGAGATACAGAATGAAA<br>GCAAGCAACCTGGCGGCAGCGCAAAAGATGCGTAAACGCAGCCTGAATGGCGAATG |
| TES-P2 | CGCAAACCGCCTCTCCCCGCAACGATCGTTGGCTGTGTTGACAATTAATCATCGGCTCGTATAATGTG<br>TGGAATTGTGAGCGCTCACAATTGGGTCATGACTGGGACACGCCAGTCATGAGAATACAGACATAAAG<br>CAAGCCAGAGATTAACGAAGCCAATTATTGAACACCCTTCGGGGTGTTTTTTTGTTTCTGGTCTACCT<br>ACTCCACCGTTGGCTTTTTTCCCTATCAGTGATAGAGATTGACATCCCTATCAGTGATAGAGATAATG<br>AGCACAGCTGTCACCGGATGTGCTTTCCGGTCTGATGAGTCCGTGAGGACGAAACAGCCTCTACAAAT<br>AATTTTGTTTAAGGGTGAATGAATTGTAGGCTTGTTATAGTTATGAACAGAGGAGACATAACATGAAC<br>AAGCCTAACCTGGCGGCAGCGCAAAAGATGCGTAAACGCAGCCTGAATGGCGAATG |
| NOR-P1 | TTAGGCGGCCGCAACGATCGTTGGCTGTGTTGACAATTAATCATCGGCTCGTATAATGTGTGGAATTG<br>TGAGCGCTCACAATTGGGTCATGACTGGGACACGCCAGTCATGAGAATACAGACATAAAGCAAGCCAG<br>AGATTAACGAAGCCAATTATTGAACACCCTTCGGGGTGTTTTTTTGTTTCTGGTCTACCACTTTTCAT<br>ACTCCCGCCATTCAGAGAAGAAACCAATTGTCCATATTGCATCAGACATTGCCGTCACTGCGTCTTTT<br>ACTGGCTCTTCTCGCTAACCAAACCGGTAACCCCGCTTATTAAAAGCATTCTGTAACAAAGCGGGACC<br>AAAGCCATGACAAAAACGCGTAACAAAAGTGTCTATAATCACGGCAGAAAAGTCCACATTGATTATTT<br>GCACGGCGTCACACTTTGCTATGCCATAGCATTTTTATCCATAAGATTAGCGGATCCGAGACCTTAC |
| NOR-P2 | TTACGGTCTCGGATCCTACCTGACGCTTTTTATCGCAACTCTCTACTGTTTCTCCATACCCGTTTTTT<br>TGGGCTAGCTACTCCACCGTTGGCTTTTTTCCCTATCAGTGATAGAGATTGACATCCCTATCAGTGAT<br>AGAGATAATGAGCACAGCTGTCACCGGATGTGCTTTCCGGTCTGATGAGTCCGTGAGGACGAAACAGC<br>CTCTACAAATAATTTTGTTTAAGGGCGTTAATCTCTGGCTTGCTTTATGTCTGTAAACAGAGGAGATA<br>CAGAATGAAAGCAAGCAACCTGGCGGCAGCGCAAAAGATGCGTAAAGCACGTACCCCGAGCCGTAGCA<br>GCATTGGTAGCCTGCGTAGTCCGCATACCCATAAAGCAATTCTGACCAGCACCATTGAAATCCTGAAA<br>GAATGTGGTTATAGCGGTCTGAGCATTGAAAGCGTTGCACGTCGTGCCGGTGCAAGCAAACCGACCAT<br>TTATCGTTGGTGGACCAATAAAGCAGCACTGATTGCCGAAGTGTATGAAAATGAAAGCGAACAGGTGC<br>GTAAATTTCCGGATCTGGGTAGCTTTAAAGCCGATCTGGATTTTCTGCTGCGTAATCTGTGGAAAGTT<br>TGGCGTGAAACCATTTGTGGTGAAGCATTTCGTTGTGTTATTGCAGAAGCACAGCTGGACCCTGCAAC<br>CCTGACCCAGCTGAAAGATCAGTTTATGGAACGTCGTCGTGAGATGCCGAAAAAACTGGTTGAAAATG<br>CCATTAGCAATGGTGAACTGCCGAAAGATACCAATCGTGAACTGCTGCTGGATATGATTTTTGGTTTT<br>TGTTGGTATCGCCTGCTGACCGAACAGCTGACCGTTGAACAGGATATTGAAGAATTTACCTTCCTGCT<br>GATTAATGGTGTTTGTCCGGGTACACAGCGTTAAGGAAACACAGAAAAAAGCCCGCACCTGACTGAGA<br>CCTTAC |
| NOR-P3 | TTACGGTCTCATGACAGTGCGGGCTTTTTTTTTCGACCAAAGGTGTCAACGTTCGACGTACGGTGGAA<br>TCTGATTCGTTACCAATTGACATGATACGAAACGTACCGTATCGTTAAGGTCAAGCTGTCACCGGATG<br>TGCTTTCCGGTCTGATGAGTCCGTGAGGACGAAACAGCCTCTACAAATAATTTTGTTTAATACTAGAG<br>AAAGAGGGGAAATACTAGATGGTGAGCAAGGGCGAGGAGCTGTTCACCGGGGTGGTGCCCATCCTGGT<br>CGAGCTGGACGGCGACGTAAACGGCCACAAGTTCAGCGTGTCCGGCGAGGGCGAGGGCGATGCCACCT<br>ACGGCAAGCTGACCCTGAAGTTCATCTGCACCACAGGCAAGCTGCCCGTGCCCTGGCCCACCCTCGTG<br>ACCACCTTCGGCTACGGCCTGCAATGCTTCGCCCGCTACCCCGACCACATGAAGCTGCACGACTTCTT<br>CAAGTCCGCCATGCCCGAAGGCTACGTCCAGGAGCGCACCATCTTCTTCAAGGACGACGGCAACTACA<br>AGACCCGCGCCGAGGTGAAGTTCGAGGGCGACACCCTGGTGAACCGCATCGAGCTGAAGGGCATCGAC<br>TTCAAGGAGGACGGCAACATCCTGGGGCACAAGCTGGAGTACAACTACAACAGCCACAACGTCTATAT<br>CATGGCCGACAAGCAGAAGAACGGCATCAAGGTGAACTTCAAGATCCGCCACAACATCGAGGACGGCA<br>GCGTGCAGCTCGCCGACCACTACCAGCAGAACACCCCAATCGGCGACGGCCCCGTGCTGCTGCCCGAC<br>AACCACTACCTTAGCTACCAGTCCGCCCTGAGCAAAGACCCCAACGAGAAGCGCGATCACATGGTCCT<br>GCTGGAGTTCGTGACCGCCGCCGGGATCACTCTCGGCATGGACGAGCTGTACAAGTAAGAATTCTTAG |

**Supplementary Table 5: Primers used in this study.**

| Primer | Template[a] | Sequence[b] |
|---|---|---|
| F_TES_1 | TES-P1 | ctaacgggggcctttttttgAACGATCGTTGGCTGTGTTG |
| R_TES_2 | TES-P1 | aaagccaacggtggagtaGGTAGACCAGAAACAAAAAAACACCC |
| F_TES_3 | TES-P2 | gtgtttttttgtttctggtctaccTACTCCACCGTTGGCTTTTTTCC |
| R_TES_4 | TES-P3 | tcctcgcccttgctcacTTTACGCATCTTTTGCGCTGC |
| R_TES_5 | pAN1720 | caacacagccaacgatcgttCAAAAAAAGGCCCCCCGTTAG |
| F_TES_6 | pAN1720 | cagcgcaaaagatgcgtaaaGTGAGCAAGGGCGAGGA |
| F_NOT_1 | pVB001 | ttcgttttggtccAGTTTACGGCTAGCTCAGTCCTAG |
| R_NOT_2 | pVB001 | gctacggctcggggtacgtgcTTTACGCATCTTTTGCGC |
| F_NOT_3 | pAN3938 | tgcgtaaagcacgtaccccgaGCCGTAGCAGCATTGGTAG |
| R_NOT_4 | pAN3938 | ttccaccgtacgtcgaaCGTTGACACCTTTGGTCG |
| F_NOT_5 | pAN4036 | caaaggtgtcaacgTTCGACGTACGGTGGAATC |
| R_NOT_6 | pAN4036 | atccggtgacagcttGACCTTAACGATACGGTACGTTTC |
| F_NOT_7 | pAN4036 | taccgtatcgttaaggtcAAGCTGTCACCGGATGTG |
| R_NOT_8 | pAN4036 | tgagctagccgtaaactGGACCAAAACGAAAAAGGC |
| F_RiboJ_Rem | pVB001, pVB002 | cgaaggtctcaGGGCGTTAATCTCTGGCTTGCTTTATG |
| R_RiboJ_Rem | pVB001, pVB002 | ggtcggtctcagcccGTGCTCATTATCTCTATCACTGATAGGGATGTC |
| F_pAN1720_EcoRI | pAN1720 | TTAGAATTCCATGGACGAGCTGTACAAG |
| R_pAN1720_NotI | pAN1720 | TTAGCGGCCGCGGCGGGAGTATGAAAAGTAAG |

a. Plasmid names prefixed with pAN are from Neilsen et al. [11].
b. Underlined lowercase sequences denote homology regions added for Gibson assembly.
